# Strength-frequency curve for micromagnetic neurostimulation through EPSPs on rat hippocampal neurons and numerical modeling of magnetic microcoil (μcoil)

**DOI:** 10.1101/2021.11.30.470598

**Authors:** Renata Saha, Sadegh Faramarzi, Robert P. Bloom, Onri J. Benally, Kai Wu, Arturo di Girolamo, Denis Tonini, Susan A. Keirstead, Walter C. Low, Theoden I. Netoff, Jian-Ping Wang

## Abstract

**Objective:** The objective of this study was to measure the effect of micromagnetic stimulation (μMS) on hippocampal neurons, by using single microcoil (μcoil) prototype, <u>Mag</u>netic <u>Pen</u> (MagPen). MagPen will be used to stimulate the CA3 region magnetically and excitatory post synaptic potential (EPSP) response measurements will be made from the CA1 region. The threshold for micromagnetic neurostimulation as a function of stimulation frequency of the current driving the μcoil will be demonstrated. Finally, the optimal stimulation frequency of the current driving the μcoil to minimize power will be estimated.

**Approach:** A biocompatible, watertight, non-corrosive prototype, MagPen was built, and customized such that it is easy to adjust the orientation of the μcoil and its distance over the hippocampal tissue in an *in vitro* recording setting. Finite element modeling (FEM) of the μcoil design was performed to estimate the spatial profiles of the magnetic flux density (in T) and the induced electric fields (in V/m). The induced electric field profiles generated at different values of current applied to the μcoil can elicit a neuron response, which was validated by numerical modeling. The modeling settings for the μcoil were replicated in experiments on rat hippocampal neurons.

**Main results:** The preferred orientation of MagPen over the Schaffer Collateral fibers was demonstrated such that they elicit a neuron response. The recorded EPSPs from CA1 region due to μMS at CA3 region were validated by applying tetrodotoxin (TTX). Application of TTX to the hippocampal slice blocked the EPSPs from μMS while after prolonged TTX washout, a partial recovery of the EPSP from μMS was observed. Finally, it was interpreted through numerical analysis that increasing frequency of the current driving the μcoil, led to a decrease in the current amplitude threshold for micromagnetic neurostimulation.

**Significance:** This work reports that micromagnetic neurostimulation can be used to evoke population EPSP responses in the CA1 region of the hippocampus. It demonstrates the strength-frequency curve for μMS and its unique features related to orientation dependence of the μcoils, spatial selectivity and stimulation threshold related to distance dependence. Finally, the challenges related to μMS experiments were studied including ways to overcome them.

## 1. Introduction

The market of neurostimulation devices and systems is estimated to reach $11 billion per year by 2026 with a compound annual growth rate (CAGR) of 12.5% [1–3]. In terms of device type, spinal cord stimulators [4–6] and deep brain stimulators [7–9] are expected lead the market, with applications to chronic pain management [10–12] and hearing loss treatments [13–15]. With the increasing rate of FDA-approved treatment options for epilepsy [16], dystonia [8], Tourette’s syndrome [7], Parkinson’s Disease [17], chronic pain [10] etc. using electrical electrodes, neuromodulation devices have a bright future. However, the existing electrical implants have their own drawbacks. They have a uniform spread of activation and must be in galvanic contact with the biological tissues. After years of implantation, due to inflammatory reactions from the neighboring tissues, the efficacy of electrical stimulation may abate due to biofouling of the electrodes [18–20]. If encapsulation occurs, electrodes may need to be replaced through a revision surgery [21,22]. While many electrodes are approved for magnetic resonance imaging (MRI), for safety concerns they are limited to low field and low resolution devices [23]. On the contrary, transcranial magnetic stimulation (TMS) is a non-invasive therapy which has been FDA-approved for depression [24] and obsessive-compulsive disorder (OCD) [25]. It uses a time-varying magnetic field that can permeate deep through the skull to stimulate the brain. This time-varying magnetic field induces an electric field in neural tissue to modulate firing rate of neurons. While there are other neuromodulation therapies, such as chemical [26], optical [27], and ultrasound [28], the neuromodulatory device presented in this work will be compared to implantable electrical electrodes and TMS therapy.

When electrical stimulation is applied through an electrode implanted on the surface, or deep in the brain, it produces very focal volumes of activation. Magnetic stimulation is applied transcranially, and while it is non-invasive it does not have the focality of implanted electrodes. The motivation for developing this device was to make a μMS device with the focality of an implantable electrode. Implantable μMS has been demonstrated previously by Bonmassar *et al*.[29] in 2012 using commercially available sub-mm sized μcoils to stimulate rabbit retinal neurons. By applying a time-varying current through a μcoil, as per Faraday’s law of electromagnetic induction, a time-varying magnetic field is generated. This magnetic field then induces an electric field in the tissue surrounding the coil, which is spatially asymmetric. The electric filed gradients generated by the μcoil may activate small, discrete populations at predictable locations relative to the coil.

Micromagnetic neurostimulation has advantage over electrical stimulation in that it doesn’t require an electrochemical interface. This allows flexibility in applying many waveforms to drive these magnetic μcoils. The application of a wide range of waveforms to drive electrical electrodes in neurostimulation are limited [30,31]. Unbalanced waveforms cause corrosion of the electrodes through irreversible redox reactions at the tissue-electrode interface. Multiple research groups have reported the fabrication of customized μcoils [32,33], of different shapes and dimensions, including planar [34,35], trapezoidal [32] or solenoidal [29,36] or V-shaped bent wires [32]. These μcoil designs have been shown to modulate different populations of neurons, in both *in vitro* and *in vivo* settings: L5 pyramidal neurons [37], intracortical neurons [32] and inferior colliculus (IC) [36] neurons. In addition, recent numerical studies have shown that these μcoils are MRI compatible [38] as they show very little to no heating around the implant area in an MRI environment. This is because there is no galvanic contact with adjacent tissues, unlike the electrical implants, thereby limiting the amount of heat generation. However, the power of operation of these μcoils is three orders of magnitude higher than DBS leads [29]. Recently, to combat this, spintronic nanodevices [39–41] having ultra-low power of operation has been theoretically proved to have therapeutic neuromodulation capability through implantable magnetic stimulation.

In this work, we have tested μcoil in rat hippocampal slice and constructed the strength-frequency curve for micromagnetic neurostimulation forecasting low power operation of these micromagnetic implants at higher frequency. To validate the EPSPs responses, we have blocked and partially reversed the response with TTX. We will also discuss the unique features of micromagnetic neurostimulation related to spatial selectivity, distance dependence and orientation dependence.

## 2. Material and methods

### 2.1. The implantable micromagnetic neurostimulation prototype

The prototype of the micromagnetic neurostimulation implant, MagPen, investigated in this work is shown in Fig. 1(a). Commercially available μcoils (Panasonic ELJ-RFR10JFB) were soldered (using solder flux and a hot air blower) at the tip of a 3 cm long pen-shaped printed circuit board (PCB). The thickness of the PCB board along the z-direction was thinned down to 0.4 mm to facilitate easier adjustment of the MagPen prototype over the hippocampal slices during *in vitro* experiments. The preferred orientation of the μcoil that would effectively stimulate neurons was unknown. Hence two different orientations of the MagPen were designed: (1) Type horizontal or Type H and (2) Type vertical or Type V. The pen-shaped tip of the Type H and Type V prototypes is 1.7 mm and 1.4 mm, respectively. The complete prototype is shown in Fig. S2(a) of Supplementary Information S2.

**Figure 1.**
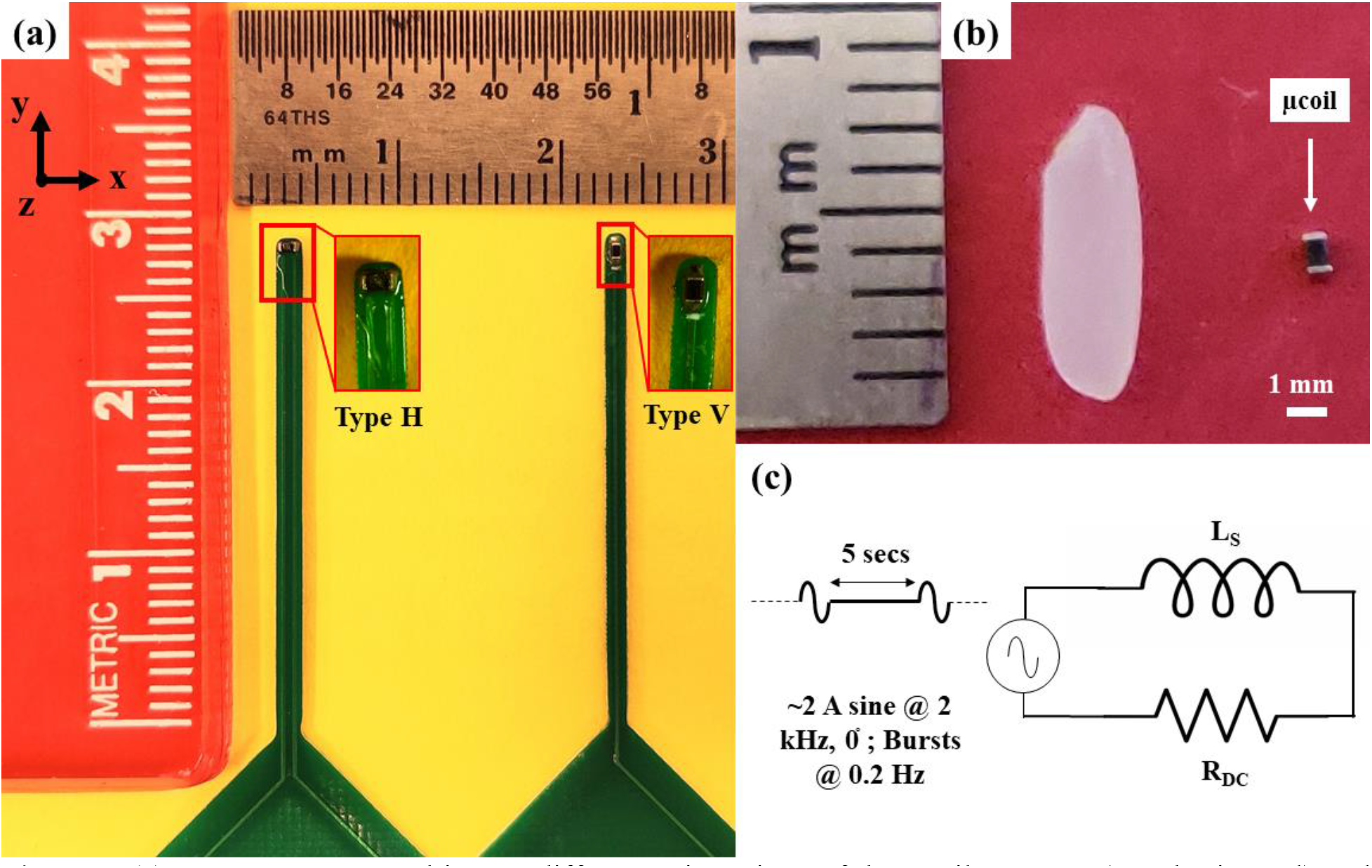
(a) MagPen prototyped in two different orientations of the μcoil: Type H (H = horizontal) and Type V (V = Vertical). The two insets show the zoomed in portion of the tips of MagPen for both Type H and Type V orientations. The μcoils are located the tip of the board and are coated with non-corrosive, biocompatible and insulation, Parylene-C. (b) The sub-mm μcoil compared to the size of a single rice grain. (c) Schematic of electrical equivalence of the μcoil as an RL circuit and diagram of the of the current waveform used drive the coil: 1-cycle of a sinusoid at 2 kHz frequency, and ~ 2 A in amplitude; each cycle of pulse is separated by 5 secs.

Both prototypes were encapsulated by 2 μm thick Parylene-C coating using the SCS LabCoater Parylene-C deposition system. The coating was used to make the MagPen prototype biocompatible, non-corrosive, and to insulate the electronics to prevent any possible neurostimulation through leakage current and capacitive coupling from these μcoils. Successful watertight and anti-leakage current coating for MagPen was ensured by measuring the impedance of one terminal of MagPen (see Fig. S2(a)) to an external electrode in artificial cerebrospinal fluid (aCSF) solution (see section 2.6). If that impedance measured above 5 MΩ, that prototype was considered to be a successful prototype for micromagnetic neurostimulation study. In the design of the micromagnetic stimulation implants, this watertight and biocompatible coating is of utmost importance to assure that any neuronal response was caused by the induced electric field only. Fig. 1(b) compares the sub-mm size of the μcoil to that of a single rice grain.

### 2.2. The electrical circuit equivalence of the μcoil

Resistance (R), inductance (L) and capacitance (C) measurements of the μcoil at 1 kHz using an LCR meter (Model no. BK Precision 889B; see Supplementary Information S1) showed that the electrical circuit equivalent of the μcoil within this frequency range is a series RL circuit (see Fig. 1(c)). Following from laws of electromagnetic induction, the inductance (L_s_) is directly proportional to the electromotive force (emf), 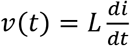, where *v*(*t*) is the emf induced in an electrical circuit (here, neurons) due to a time-varying magnetic flux density; *v*(*t*) contributes directly to the induced electric field which is used to stimulate the neurons and 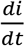 is the time-varying current through the inductor or μcoil. The resistance (R_DC_) contributes to heat dissipation from the electrical circuit. Future designs of the micromagnetic neurostimulation implants will be focused on further reducing the resistance because lower resistance coils will generate less heat and for the same magnetic field.

### 2.3. The μcoil driving circuitry

To test the μcoils in brain slice, they were powered by 1-cycle of sinusoidal current of amplitudes 2 A, frequency 2 kHz, phase 0° and offset at 0 mV. Each cycle of pulse is separated by a duration of 5 secs (see Fig. 1(c)). The 5 sec pause between the stimulation pulses was to assure synaptic plasticity effects were not induced in the synaptic coupling between the hippocampal neurons [42,43]. The current waveform was generated using a function generator (model no. Tektronix AFG2021) and amplified using a MOSFET class-D amplifier of 70 kHz bandwidth (model no. Pyramid PB717X). Fig. 2(d) – left shows the signal flow chain to power the μcoils for EPSP recording studies in this work.

**Figure 2.**
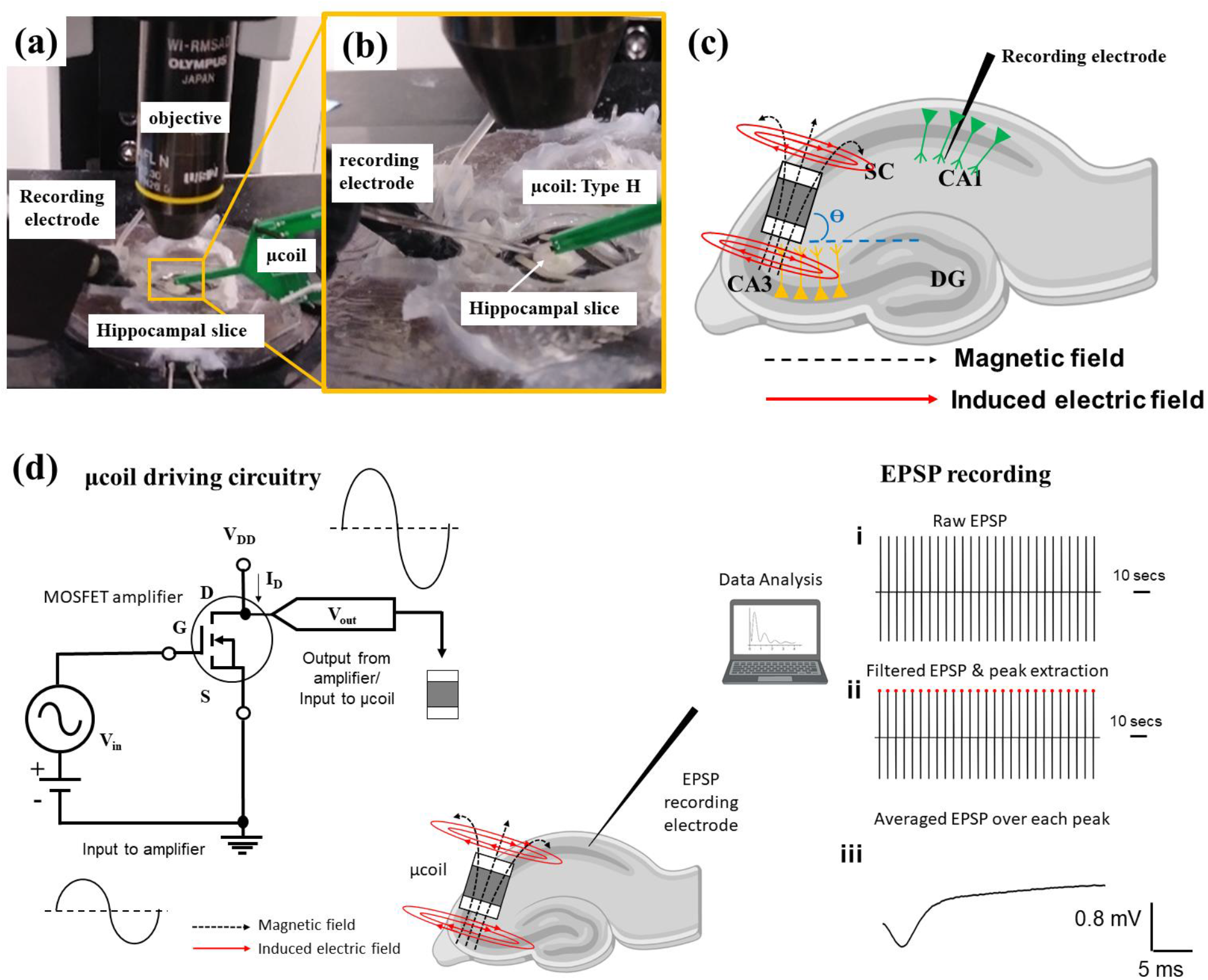
*in vitro* experimental set-up. (a) MagPen, Type-H orientated over the rat hippocampal slice for electrophysiological recordings. (b) Magnification of the chamber containing the hippocampal slice and μcoil in the preferred orientation, Type H for *in vitro* stimulation. (c) Orientation of the μcoil over Schaffer Collateral fibers which induces excitatory post synaptic potential (EPSP) from the CA1 neurons. (DC: Dentate Gyrus; CA3: Cornu Ammonis 3; SC: Schaffer Collateral; CA1: Cornu Ammonis 1). (d) μcoil driving circuitry (left): a function generator generating sinusoidal waveforms (V_in_) is amplified using a MOSFET amplifier with a user-controlled gain. The amplifier output (V_out_) is used to drive the μcoil. EPSP recording (right): The raw EPSPs in (i) were recorded from the hippocampal slices. EPSPs were then analyzed through a customized algorithm MATLAB script to remove noise and estimate peak amplitude, an example of an EPSP is shown in (ii). Finally, EPSPs from approximately 20-25 trials were averaged to estimate the EPSP amplitude from CA1 neurons due to μMS over the SC fibers as in (iii).

### 2.4. Electromagnetic modeling

Measuring the magnetic flux density and the induced electric field from these sub-mm sized μcoils is difficult experimentally difficult requiring custom-made miniature-sized pick-up coils [44] and close proximity of the μcoils. Reliable numerical modeling using finite element method (FEM)-based calculations to study induced currents in neural tissues have been previously reported to characterize both the electrical and magnetic fields [45–47]. Therefore, we conducted a FEM modeling study of the μcoil using ANSYS-Maxwell [48] eddy current solver (ANSYS, Canonsburg, PA, United States) which solves a modified version of the T-Ω formulation of the Maxwell’s equations [49]. The ceramic core μcoil dimensions, tissue slab parameters, boundary conditions and the high-resolution tetrahedral mesh size used are detailed in Table 1. Simulations were done using the Minnesota Supercomputing Institute (MSI) at the University of Minnesota (8 cores of Intel Haswell E5-2680v3 CPU, 64×8=512 GB RAM and 1 Nvidia Tesla K20 GPU). The induced electric field values were then exported to be analyzed using a customized code written in MATLAB (The Mathworks, Inc., Natick, MA, USA).

**Table 1.** Electromagnetic modeling parameters

| Parameter description | Value |
| --- | --- |
| μcoil dimension (L × W × H) | 1 mm × 600 μm × 500 μm |
| no. of turns (N) | 21 |
| Wire diameter | 7 $\mu\text{m}$ |
| Tissue dimension (L × W × H) | 4 mm × 4 mm × 300 $\mu\text{m}$ |
| Conductivity of tissue ($\sigma$) | 0.13 S/m |
| Air dimension (L × W × H) | 10 mm × 10 mm × 4 mm |
| Energy error (user-specified) | 1 % |
| Final solution no. of mesh elements | 410,000 |
| Adaptive passes (converged) | 6 |

### 2.5. Modeling using NEURON

To simulate the effects of electric field induced by the magnetic field generated by the MagPen, we modified a model of a layer 5 (L5) pyramidal neuron developed by Pashut *et al*. [50]. In their model the spatial component of the induced electric filed generated by a transcranial magnetic stimulation (TMS) coil is projected into a 4 mm × 4 mm array in MATLAB. Then simulations of the pyramidal neuron in the NEURON package [51,52] were performed at different positions of the neuron relative to the center of the coil (see section 3.2). We modified Pashut’s NEURON [50] by replacing the array of the induced electric with ours generated from the FEM-model of the μcoils from ANSYS-Maxwell (see section 2.4). Simulations of the time varying waveform at 2 A amplitude sinusoidal current at 2 kHz through the MagPen were carried out. The membrane potential at the soma was then measured and the volume of activation around the MagPen estimated.

### 2.6. The hippocampal slice preparation and EPSP recording

All brain slicing experiments (see Fig. 2(a)) were done in accordance with a protocol approved by the University of Minnesota Institutional Animal Care and Use Committee (IACUC). Brain slices were prepared at 300 μm - 400 μm thickness (see Fig. 2(b) & (c)) from 14- to 21-day-old Long-Evans (L/E) rats using a PELCO easiSlicer™ Vibratory Tissue Slicer (Ted Pella, Inc.). Slices were immediately incubated in standard aCSF with a composition (in mM) of 124 NaCl, 2 KCl, 2 MgSO4, 1.25 NaH2PO4, 2 CaCl2, 26 NaHCO3, and 10 D-glucose [53] at 33 °C for at least 1 hour. The aCSF solution was oxygenated with a 95% O2 and 5% CO2 for the duration of the experiment. EPSPs were measured in the CA1 at 10 kHz using a glass capillary microelectrode filled with aCSF (3-4 MΩ) with a MultiClamp 700B Microelectrode Amplifier (Molecular Devices), Fig. 2(d).

## 3. Results

### 3.1. Preferable μcoil orientation using modeling, position of neuron and validation using experiments

As shown in Fig. 1(a), two prototypes for MagPen have been prepared in a horizontal (Type H) and vertical (Type V) orientations. The ideal location and orientation of the coil activate the Schaffer Collateral fibers in the hippocampal slice (including the pyramidal cell layers) are shown in Fig. S2 (b) & (c) in Supplementary Information S2.

In Fig. 3(a) - i, ii & iii, the spatial heat maps of the induced electric field from Type H orientation of the μcoil over a biological tissue at 300 μm distance between tissue and the μcoil are shown. In Fig. 3(b) - i, ii & iii, the spatial heat maps of the induced electric field from Type V orientation of the μcoil over a biological tissue at 300 μm distance between tissue and the μcoil are shown. In both Fig. 3(a) & (b), the μcoil is driven by a sinusoidal current of amplitude 2 A and frequency 2 kHz. The size of the neural tissue was 4 mm × 4 mm × 300 μm. The position of the modeled neuron is shown in Fig. 3(a) & (b)- i, ii & iii. To study the effects of micromagnetic neurostimulation, only the in-plane components, E_x_ and E_y_ of the induced electric field (E), were used in the model. Modeling the membrane potential from the neurons situated at a specified location (see Fig. 3(a) & (b)) for both Type H and Type V oriented μcoils, action potentials were detected for the neuron being stimulated by Type H orientation only (see Fig. 3(a) - iv). For Type V orientation, the Ex-component of the induced electric field is negligible (see Fig. 3(b) - ii) failing to induce an action potential in the neuron (see Fig. 3(b) - iv).

**Figure 3.**
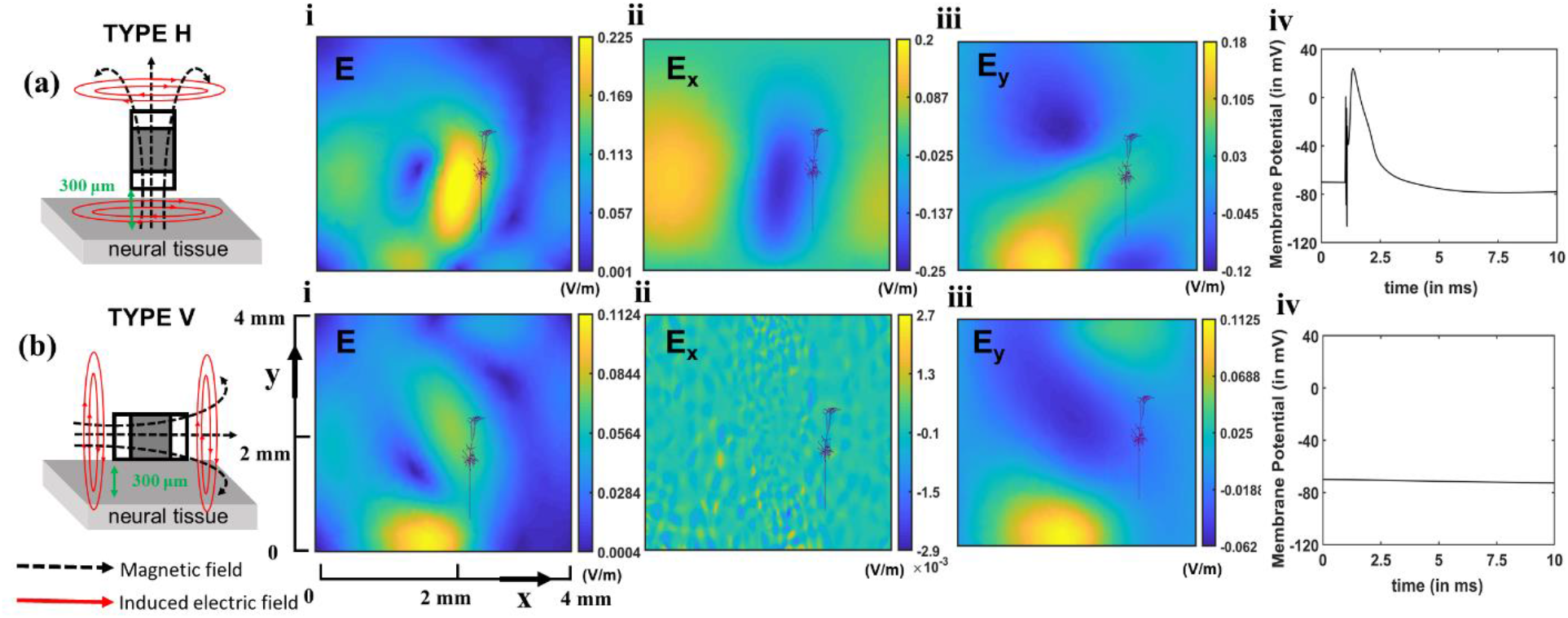
Modeling results corresponding to ‘ideal’ orientation of MagPen case for successful micromagnetic neurostimulation. The μcoil was driven by a sinusoidal current of amplitude 2 A at a frequency of 2 kHz. The approximate position of the modeled neuron with respect to the tissue has been shown. (a) MagPen, Type H: Spatial heat maps for the induced electric fields calculated for a tissue dimension of 4 mm × 4 mm × 300 μm, at 300 μm from the μcoil surface for (i) E (in V/m) (ii) E_x_ in V/m (iii) E_y_ in V/m (iv) membrane potential in the neuron; (b) MagPen, Type V: Spatial heat maps calculated for a tissue dimension of 4 mm × 4 mm × 300 μm at 300 μm from the μcoil surface for (i) E (in V/m) (ii) E_x_ in V/m (iii) E_y_ in V/m (iv) membrane potential in the neuron. Type H orientation in (a) elicited an action potential, but Type V orientation in (b) did not elicit an action potential. x and y in (b-i) denote the spatial axes coordinates in the neural tissue. It is same for all the spatial heatmaps.

In an ideal scenario, as demonstrated in Type H orientation in Fig. 3(a), the angle between the μcoil and the neural tissue were required to be 90°. However, in the *in vitro* experiments it was difficult to orient the Type H probe at an angle less than 90°. This was an unavoidable experimental challenge that created a difference between ‘ideal’ and ‘real’ orientation of Type H MagPen over the Schaffer collateral. In this *in vitro* experiment we recorded EPSPs from the CA1 region of rat hippocampal slices by stimulating the SC fibers using the MagPen. For the experimental set-up, Type H orientation (see Fig. 2(a) & (b)) was found to elicit population EPSPs in neurons in the CA1.This observation was in corroboration with our FEMbased modeling studies done on ANSYS-Maxwell.

### 3.2. Spatial selectivity and distance dependence in micromagnetic neurostimulation

For the Type H orientation of the MagPen, there were two unique features specific to micromagnetic neurostimulation observed. In Fig. 4(a)-i for the Type H orientation of the μcoil at a distance of 300 μm from the neural tissue, activated approximately 2.672 mm^2^ of tissue. The most sensitive area of the tissue is marked by the ellipsoid in dotted black line where the μcoil position was at the center of the tissue marked by solid red line (see Fig. 4(a)-ii). On rotating the μcoil along any direction a completely new tissue area can be activated.

**Figure 4.**
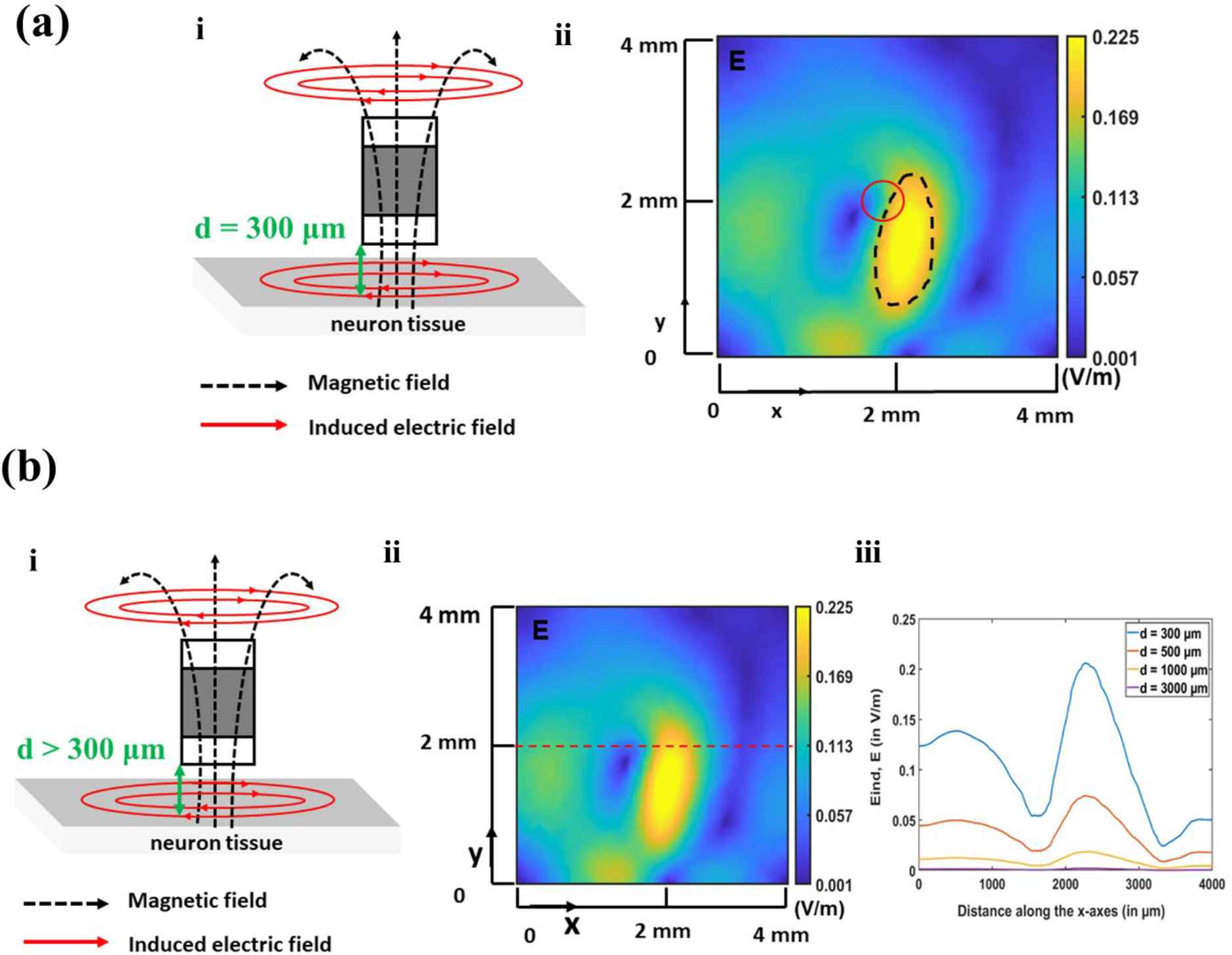
Spatial selectivity and distance dependence of MagPen. (a) Spatial selectivity of MagPen, Type H. (i) For a distance, d = 300 μm between the μcoil and the neural tissue, (ii) only 16.7 % of the tissue area (denoted by the dotted black ellipsoid) has the maximum probability of getting stimulated. The location of the μcoil denoted by red solid circle. This implies that μMS is spatially selective. (b) (i) For a distance, d > 300 μm between the Type H oriented μcoil and the neural tissue, (ii) the induced electric field value was measured along x-axes, along the dotted red line. (iii) As the distance between the μcoil and the neural tissue was increased, the induced electric field value decreased drastically. Our NEURON model showed that beyond 300 μm distance between the μcoil and the neural tissue, no action potential was observed. The μcoil was driven by a 2 A sinusoidal current at 2 kHz frequency. All measurements were made by varying the distance between the μcoil and the neural tissue of dimension 4 mm × 4 mm × 300 μm. x and y in (a-ii) & (b-ii) denote the spatial axes coordinates in the neural tissue. It is same for all the spatial heatmaps.

When the same Type H MagPen orientation was modeled, if the distance between the μcoil and neural tissue was greater than 300 μm, we observed a significant attenuation of the induced electric field (Fig. 4b). Validating the neurostimulation capability of this μcoil under this condition, the optimum distance for activation of a neural tissue was found to be 300 μm. Beyond this distance, no activation of the neuron was observed. Hence, in experiments on animal model, manipulating the correct distance between the μcoil and the neural tissue is extremely critical. The range of attenuation can be estimated from Fig. 4(b)-iii which is measured for distances between μcoil and tissue, 300 μm, 500 μm, 1000 μm, and 3000 μm along the x-axes of the neural tissue intersecting the y-axes at the center (see Fig. 4(b)-ii). A detailed demonstration of the attenuation of induced electric field (E) and its in-plane components E_x_ and E_y_ for distances between the tissue and μcoil above and below 300 μm has been demonstrated in Supplementary Information S3.

In the simulation studies for both Fig. 4 (a) & (b), the μcoil was powered by 2 A sinusoidal current in 2 kHz frequency. The features of spatial selectivity and distance dependence of the μcoil are unique features for micromagnetic neurostimulation, which can potentially offer targeted, precise, highly focal, and spatially tunable neurostimulation capability.

### 3.3. Strength-frequency curve for micromagnetic neurostimulation

We investigate the strength-frequency relationship which is the micromagnetic neurostimulation equivalent of the strength-duration relationship described for electrical stimulation. The induced electric field equation, which follows from Faraday’s Law of electromagnetic induction, can be written as:

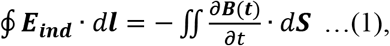

where, ***B*** is the magnetic flux density generated from the μcoil, ***E_ind_*** is the induced electric field that will stimulate the neurons, ***l*** and ***S*** are the contour and the surface area of the neural tissue.

In both our numerical calculations as well as in experiments we have used a sinusoidal current waveform to drive the μcoil. Therefore, the current waveform can be implemented as: *i*(*t*) = *I*_0_*sin* (2*πft*), where, *I*_0_ is the amplitude of the current, *f* is the frequency of the current and *t* is the time instant. The magnetic flux density is represented by: 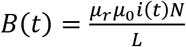 where, *μ_r_* is the relative permeability of the medium, *μ*_0_ is the vacuum permeability, *N* is the number of turns of the μcoil and *L* is the length of the μcoil. The spatial heatmap for the magnetic flux density (B_x,y,z_) is shown in Fig. S6 of Supplementary Information S6, whereas the temporal component of the magnetic flux density is equivalent to that of the current driving the μcoils.

On substituting the values for *i*(*t*) and *B*(*t*) in equation (1) followed by a simple differentiation, a modified version of induced electric field is obtained:

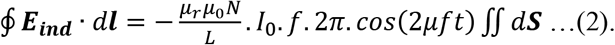

Observing equation (2), we see that the parameters, *μ_r_, μ*_0_, *I* and *S* are all dependent on the neural tissue while, *N* and *L* are dependent on the μcoil. Both these groups of parameters cannot be altered and are specific to the neural tissue that we are trying to stimulate and the μcoil model that we are trying to use. There are only two parameters that we can alter, *I*_0_ and *f*. *E_ind_* is directly proportional to both *I*_0_ and *f*. Equation (2) also demonstrates that the temporal component of the induced electric field will be a time derivative of the current driving the μcoils.

The working window obtained from numerical modeling on ANSYS-Maxwell in Fig. 4 shows that keeping the amplitude of the current constant at 2 A, increasing the frequency of the current driving the μcoil from 500 Hz to 5 kHz, the induced electric field (E) amplitude also increases. This is evident from the spatial heat maps for E, E_x_ and E_y_ measured for a biological tissue of dimension 4 mm × 4 mm × 300 μm. The distance between the μcoil and the neural tissue has been 300 μm in all cases. This working window in Fig. 5 implies that at higher frequency, the amplitude of the current required to drive the μcoils to trigger an action potential will be less. This means, at higher frequency, the μcoils will have low power consumption and this in terms of micromagnetic neurostimulation imply significant reduction of thermal effects on tissues.

**Figure 5.**
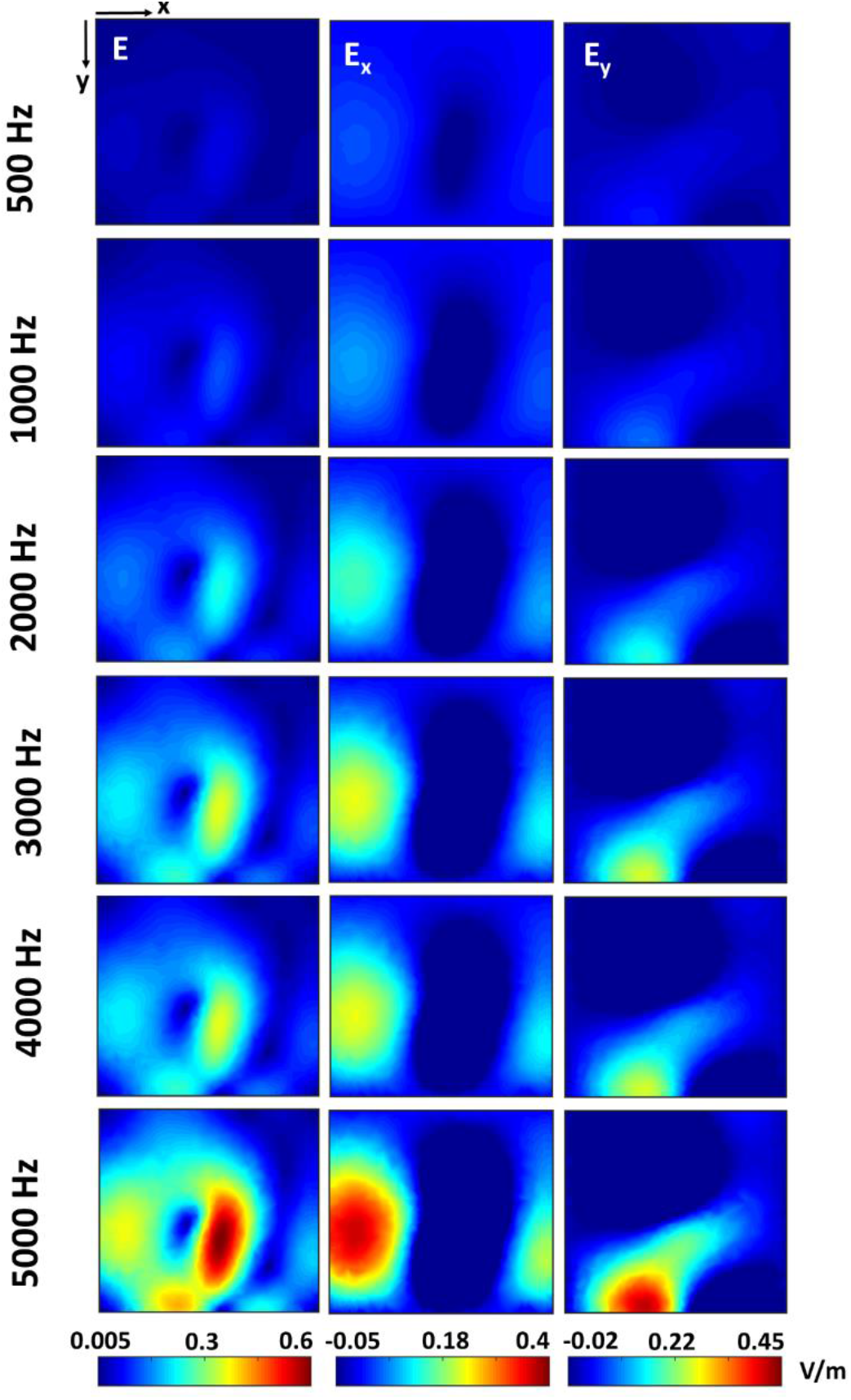
Effects of varying stimulation frequency on electric fields. Working window of the spatial heat maps of the x-component (E_x_ in V/m) and the y-component (E_y_ in V/m) of the induced electric (E in V/m) on neural tissue of dimension 4 mm × 4 mm × 300 μm at different frequency of the 2 A of sinusoidal current driving the μcoil. With increasing frequency, the induced electric field value increased. From an experimental point-of-view, this suggests that at higher frequencies, the current amplitude required to stimulate a neuron should decrease. x and y denote the spatial co-ordinate axes directions of the tissue.

Through subsequent numerical calculations on ANSYS-Maxwell [48] and NEURON [51] we plotted the ‘strength-frequency’ curve for micromagnetic neurostimulation specific for this μcoil model in Fig. 6. The heat maps at different sinusoidal current amplitudes and frequencies obtained from ANSYS-Maxwell (eddy current solver), were input to the NEURON model. Then it was calculated at which current amplitude and frequency combination, the neuron elicits an action potential. Each of the points on the graph in Fig. 6 represent the current amplitude and frequency combination at which an action potential was observed. This implies that to elicit an action potential, at lower frequencies, we require a higher current amplitude and at higher frequency we require a lower current amplitude. However, in experiments (see section 3.4), the current amplitude required to stimulate a neuron at a certain frequency was found to be slightly greater because of some challenges discussed in section 4 in details.

**Figure 6.**
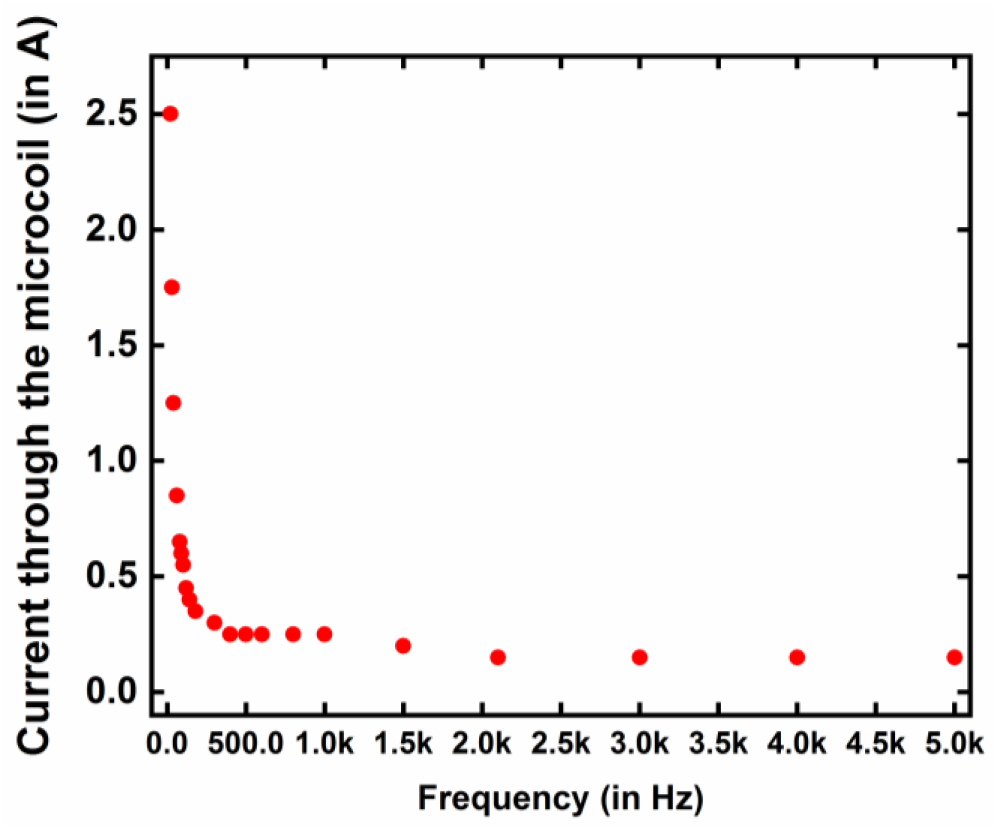
The strength-frequency curve for MagPen established through ANSYS-Maxwell and NEURON modeling.

### 3.4. Validation of biological neuronal responses using TTX and TTX washout experiments

In experiments, due to experimental limitations, it was not possible to place the μcoil in the ‘ ideal’ orientation, as discussed later (see Fig. 2(c) & Fig. S2(a), Supplementary Information S2). In this work, as shown in Fig. 2(d) *in vitro* EPSP recordings from CA1 region of the rat hippocampal slices were measured following stimulation of the Schaffer collaterals with the MagPen. The raw EPSPs were recorded, filtered, and averaged over at least 20-25 trials. Due to experimental challenges discussed in section 4 concerning spatial selectivity, distance dependence, and correct orientation of the MagPen prototype, and stimulus artifacts, EPSPs were validated by blocking the neural response using the voltage-gated Na channel blocker TTX [54]. The steps have been pictorially summarized in Supplementary Information S4. Because TTX blocks voltage gated Na channels, it blocks action potentials in Schaffer collaterals generated by the MagPen and the resulting EPSPs in CA1 are eliminated. After washing out the TTX we would expect a complete, or at least a partial, recovery of the EPSP in response to magnetic stimulation. Fig. 7(a)-i shows the EPSP recorded from CA1by MagPen Type H stimulation on the Schaffer collaterals. Without altering the position of the MagPen and the EPSP recording electrode, the slice was superfused with artificial CSF containing 1 μM of TTX for 35 – 40 mins. At this time, the magnetic stimulation was applied, and the recording is shown in Fig. 7(a)-ii. Finally, response after 10 minute TTX washout is shown in Fig. 7(a)-iii.

**Figure 7.**
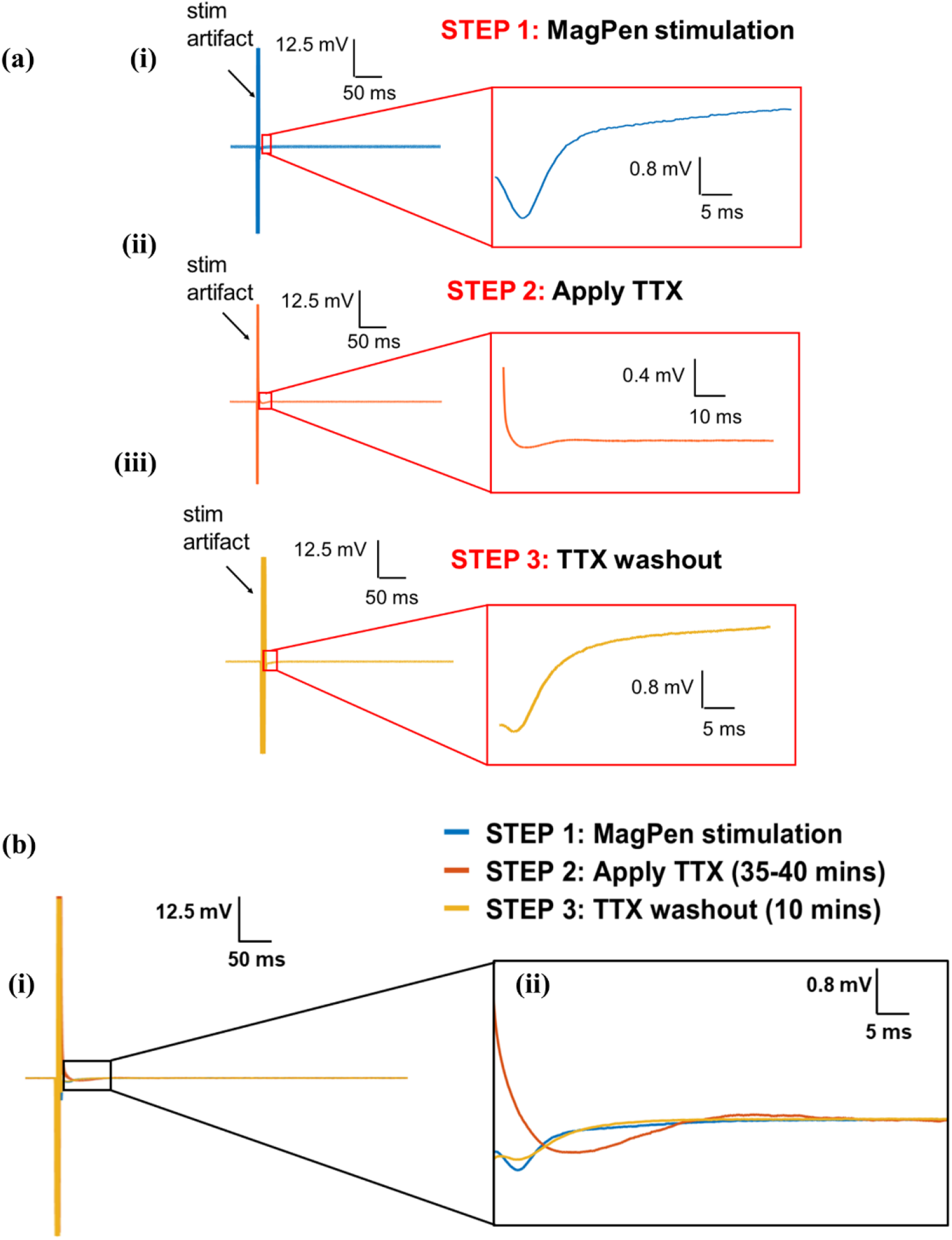
Validating whether the EPSP recording from MagPen stimulation is a biological response. In all cases, MagPen powered by 2 A @ 2 kHz sinusoidal pulse @ 5 secs. The MagPen stimulation was applied at the CA3 region and the EPSP was recorded at the CA1 region. The EPSP recorded from: (a)-(i) STEP 1 – MagPen stimulation only: Main EPSP of the averaged signal is zoomed out by omitting the stimulation artifact. (ii) STEP 2 – Apply TTX (35 – 40 mins): To the same hippocampal slice, 1 μM of TTX was perfused at a rate of 0.8 – 1 ml/min for 35 – 40 mins. Main EPSP of the averaged signal is zoomed out by omitting the stimulation artifact. (iii) STEP 3 – TTX washout (10 mins): To the same hippocampal slice, TTX was washed out by perfusing aCSF solution at a rate of 0.8 – 1 ml/min for 10 mins. Main EPSP of the averaged signal is zoomed out by omitting the stimulation artifact. (b) Overlapped EPSPs obtained from the 3 steps in (a). (b)-(i) Averaged EPSPs overlapped in STEP 1, STEP 2 and STEP 3. (ii) The average signals zoomed in, obtained by removal of the stimulation artifact. The removal of the trough from STEP 1 (trace in blue) in STEP 2 (trace in orange) implies that TTX successfully blocked the neuron response in STEP 1. In addition, the signal observed in STEP 1 is indeed biological. Successful TTX washout in STEP 3 can be concluded from the partial reappearance of the trough in STEP 3 (trace in yellow). This further strengthens the fact that the trace in blue obtained from magnetic stimulation only is indeed biological. This was observed in 4 out of 6 trials of this 3-step experiment.

The superposition of the EPSPs measured before, during TTX and after washout are shown in Fig 7(b). It can be seen that the EPSP (blue trace, from Fig. 7(a)-i) disappeared after application of TTX (orange trace, from Fig 7(a)-ii). This suggests that the signal induced by magnetic was dependent on neuronal action potentials. After washout of TTX the trough observed before TTX was almost completely recovered (yellow trace, from Fig. 7(a)-iii). These results support the conclusion that the signal observed is an EPSP and not an electronic artifact.

## 4. Discussion

Micromagnetic neurostimulation is still in its infancy with limited biologically meaningful results. However, the involvement of several research groups worldwide in this line of research has made it increasingly popular. Three unique features of this technology that had been previously identified, have been quantified in this work: (1) orientation dependence, (2) spatial selectivity and, (3) depth dependence.

First, with respect to orientation dependence, there has been enough controversy in the literature regarding what orientation is the most suitable to observe elicitation of a neuron response, in both *in vitro* and *in vivo* settings. Bonmassar *et al*. [29], Lee *et al*. [55] and Osanai *et al*. [56] have performed independent calculations and experiments to arrive at a necessary conclusion. While Bonmassar *et al*. [29] in their very first works mentions how both orientations can elicit a neuron response that can be recorded using patch clamp set-up, Lee *et al*. [55] mentions that Type H orientation over the pyramidal neurons is second most preferable option to observe a neuron response with Type V orientation being the most preferable. On the contrary, Osanai *et al*.[56] in their *in vivo* study on cortical neurons mentions how they observed local field potentials (LFP) using Type H orientation only on the cortex up to a significant depth of 950 μm. In this work, we corroborate with Osanai *et al.’s* [56] work to conclude that Type H orientation is the preferable option to elicit a neuron response (see section 3.1 & 3.4). However, further research studies are encouraged to establish this point because the performance of micromagnetic neurostimulation can vary between different population of neurons in both *in vitro* and *in vivo* studies. It might also be highly dependent on the orientation of the neurons in the experimental study settings. Hence, preliminary studies involving this technology should consider a well investigated neuron model in experiments (with known electric field threshold and frequency) as in this work we used the well oriented Schaffer Collateral fibers of the hippocampal slice as the experimental model.

Second, the quantified spatial selectivity of these μcoils demonstrated in Fig. 4(a) for this μcoil model is reported for the first time in the literature of micromagnetic neurostimulation. We demonstrated that a particular position of this μcoil model driven by a 2 A sinusoidal current waveform at 2 kHz frequency activates 2.672 mm^2^ of the tissue area can be stimulated at a time. Rotating the μcoil along any direction – clockwise or anticlockwise - will activate a completely different 2.672 sq mm of the tissue area. Hence, during the *in vitro* experiments to record EPSPs from hippocampal slices, care had to be taken that at a time one does not accidentally rotate the MagPen prototype in the micromanipulator. This fact also justifies why we could not validate the EPSPs from 2 trials out of 6 trials in the 3-step TTX-TTX washout experiments demonstrated in Section 3.4 and Supplementary Information S5. The amount of tissue area that is expected to be activated will vary with the dimension of the μcoil and the distance between the μcoil and the tissue.

Third, this work numerically established the distance dependence of micromagnetic stimulation and the threshold distance for this μcoil model. The numerical simulations suggested that it was essential to maintain the distance between the μcoil and the tissue well below 300 μm. Beyond 300 μm our modeling results from the NEURON model showed no neurostimulation effect (see Supplementary Information S3). Therefore, this distance dependent feature for micromagnetic neurostimulation promises targeted and local activation of neurons. As much as this feature is the ardent need for next generation neuromodulation devices, this also makes EPSP recordings from micromagnetic neurostimulation extremely challenging. During our *in vitro* experiments, it was essential to keep the distance between μcoil and the neural tissue well below 300 μm. This also means we must control the thickness of the Parylene-C coating on the μcoil. While using the micromanipulator to adjust the distance between the μcoil and the hippocampal tissue, it is critical to adjust the distance between the μcoil and the tissue below 300 μm.

In Section 3.3, the strength-frequency curve for this μcoil model has been established using ANSYS-Maxwell and NEURON modeling. In experiments (discussed in section 3.4) the threshold current required to drive the μcoil such that we can stimulate the CA3 neurons (marked in yellow in Fig. 2(c)) at 2 kHz frequency is ~ 2 A. However, the strength-frequency curve in Fig. 6 reads ~ 0.25 A. The reason behind the higher threshold in experiments can be justified by the challenges faced during the *in vitro* experimental set-up in Fig. 2(a) related to orientation dependence, spatial selectivity and distance sensitivity. If the μcoil model is changed, the trend might still be the same, but the numerical values will vary.

We chose hippocampal slices because the well aligned axons from CA3 projecting to the CA1 region through the Schaffer collateral (see Fig. 2(c)) make this a common preparation for electrophysiological experiments. Furthermore, seizures have been induced in hippocampus in that past, making it a suitable subject for studying epilepsy [57]. Although this technology supports spatially selective neurostimulation, we did not face difficulty in deciding which set of neurons to record from. As we chose to study the CA3-CA1 hippocampal synaptic pathway, we kept the recording electrode fixed in the CA1 region. On stimulating the CA3 region using MagPen prototype, one could record EPSPs from the synaptically activated CA1 neurons (marked in green in Fig. 2(c)) if the Type H prototype was oriented at a distance below 300 μm between the μcoil and the tissue.

Reproducibility for *in vitro* experiments related to micromagnetic neurostimulation is a pressing issue due to several challenges related to electromagnetic interference (EMI), spatial selectivity and distance dependence. Nevertheless, with tedious grounding efforts and the correct adjustment of MagPen prototype orientation over the hippocampal tissue we have been able to obtain biological EPSPs from micromagnetic stimulation. The EPSPs could be suppressed by application of TTX in 4 out of 6 trials with an average of 20 % drop in the EPSP peak (see Supplementary Information S5). We also observed partial return of EPSPs on TTX washout with a success rate in 2 out of 3 trials of successful suppression on TTX application (see Supplementary Information S5).

There is ample research scope in this interesting yet challenging field of research. One can envision development of a portable, robust, and user-friendly waveform generator system [58] specific to drive these μcoils such that micromagnetic neurostimulation can be translated to be studied in a clinical setting. Besides, the development of custom-fabricated μcoils [35] in order to lower the power of operation and make the induced electric field even more spatially focused will be an interesting way to proceed with development in this field. In addition, it would be interesting to know how arrays of μcoils [34] can spatially activate different populations of neurons. The high distance sensitivity with which these magnetic μcoils can stimulate the neurons has been studied. At the same time, it has been predicted how miniscule these μcoils can be fabricated to facilitate cellular level neurostimulation [34,35]. Hence, for this technology of micromagnetic neurostimulation to reach clinical settings, it requires advancement in neurosurgical techniques too. Finally, since these μcoils use magnetic field to stimulate the neurons, there is ample scope of using a wide variety of waveforms to stimulate the neurons unlike electrical electrodes where due to safety limitations, one cannot use all types of waveforms for neurostimulation. Advancement in μcoil designs promises implantable neuromodulation that may complement or be an alternative to electrical stimulation.

## 5. Conclusion

This work tests a novel micromagnetic coil for neuromodulation in a hippocampal slice. Only one orientation of the μcoil, Type H-MagPen prototype, could successfully activate neurons; hence they demonstrate orientation-specific activation. For the first time we have quantified that a tissue area of 2.672 mm^2^ can be activated at a time for a particular location of this μcoil model. This contributes to spatially selective stimulation capability of the μcoils. We investigated this using numerical simulations and corroborated through *in vitro* EPSP recording experiments that this μcoil at 2 A sinusoidal current of frequency 2 kHz cannot activate neurons beyond a μcoil-tissue distance of 300 μm. The EPSPs averaged over 20-25 trials from the micromagnetic stimulation were validated to be successful biological responses using a previously reported TTX. On application of TTX, the EPSPs dropped on an average of 20% (calculated over 4 successful trials out of 6). On TTX washout, partial return of the EPSPs was observed; the difference between original EPSP peak and the peak after TTX washout is 0.1 % on an average. Finally, we demonstrated the strength-frequency curve specific for MagPen which showed that at higher frequency of the driving current, lower current amplitudes are required to activate the neurons.

## Notes

The authors declare no conflict of interest.

## Acknowledgements

This study was financially supported by the Minnesota Partnership for Biotechnology and Medical Genomics under award number ML2020. Chap 64. Art I, Sec11on 4. R.S. acknowledge the 3-year College of Science and Engineering (CSE) Fellowship awarded by University of Minnesota, Twin Cities. The authors would also like to thank useful discussions Dr. Winfried A. Raabe, M.D. from the Department of Neurosurgery; Kendall H. Lee, M.D., PhD, Charles D. Blaha, PhD and Yoonbae Oh, PhD from Mayo Clinic, Rochester, MN. Portions of this work were conducted in the Minnesota Nano Center (MNC), which is supported by the National Science Foundation through the National Nano Coordinated Infrastructure Network (NNCI) under Award Number ECCS-2025124. J.P.W also thanks the Robert Hartmann Endowed chair support.

