## Supplementary Information for "Strength-frequency curve for micromagnetic neurostimulation through EPSPs on rat hippocampal neurons and numerical modeling of magnetic microcoil (μcoil)"

### S1. RLC measurements of the µcoil affecting the choice for µcoil in µMS

Resistance (R), inductance (L) and capacitance (C) measurements of the µcoil at 1 kHz from LCR meter (Model no. BK Precision 889B), (see Table 1) showed that the electrical circuit equivalent of the µcoil within this frequency range is a series RL circuit (see Fig. 1(c)). Reactance due to capacitance is given by $\frac{1}{wC}$, while reactance due to inductance is given by $wL$; where $w = 2\pi f$ and $f$ is the frequency at which the measurement is made, here 1 kHz. Therefore, higher the capacitance, lower is the impedance, implying less resistance to the flow of current. But higher the inductance, higher is the impedance, implying greater resistance to the flow of the current.

Ohm's Law, $V = iR$, where V = voltage, i = current and R = resistance, states that current always travel through the path of lowest resistance in an electrical circuit. From the RLC parameters of the µcoil measured in Table 1 (the measurements varied with an error margin of ±0.0001% between different µcoils due to difference in soldering conditions), the series capacitance ($C_s$) being small offer low resistance to the flow of the current. While the parallel inductance ($L_p$) being high and the parallel capacitance ($C_p$) being low, both offered higher resistance to the flow of current. Therefore, whatever current was applied to the µcoil travelled through the path of lowest impedance, the resistance, and the series inductance ($L_s$).

The µcoil model no. used in this work (Panasonic ELJ-RFR10JFB) is currently an obsolete component in the market. The reason for the choice of this specific µcoil model has been demonstrated in this section, Supplementary Information S1. However, one can always find better alternatives in the market by looking for µcoils whose electrical circuit components reduce to a series RL circuit with an even lower resistance. Custom designing µcoils to facilitate micromagnetic neurostimulation is also a viable option.

Table 1. RLC measurements of the µcoil

| **Parameters** | **Value** |
|---|---|
| Type of coil | solenoid |
| No. of turns (N) | 21 |
| Resistance R | 3.6678 – 4.4 Ω |
| Series Inductance $L_s$@1kHz | 0.654 – 0.71 µH |
| Series Capacitance $C_s$@1kHz | 25 mF |
| Parallel Inductance $L_p$@1kHz | 4 mH |
| Parallel Capacitance $C_p$@1kHz | 76.83 nF |

**S2. 'Ideal' orientations of Type H and Type V prototypes over the Schaffer Collateral fibers of the rat hippocampal slice**

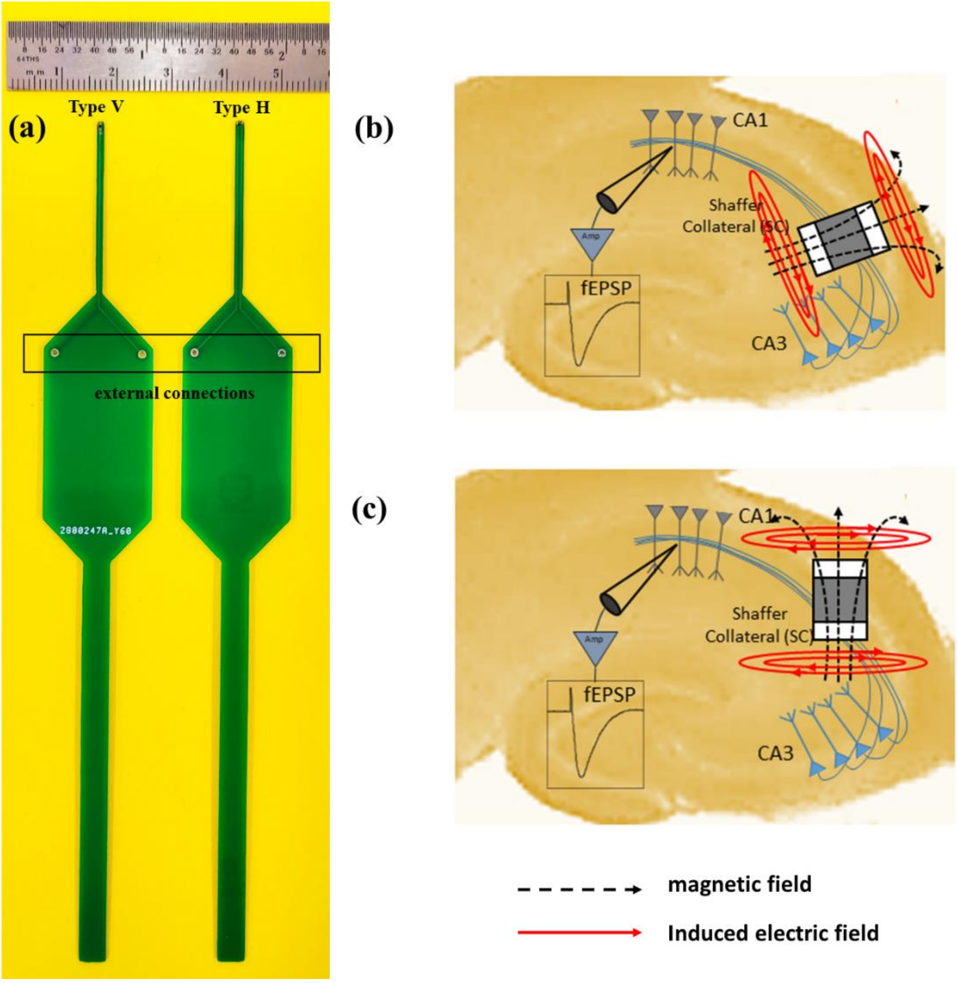


Figure S2. (a) The complete image of the MagPen prototype, both Type V and Type H. 'Ideal' orientation of (b) Type V and (c) Type H over the Schaffer Collateral fibers to study the micromagnetic neurostimulation performance of MagPen.

Fig. S2(a) shows the external terminal connection for the MagPen prototype. From those external connection pins, wires were soldered out to create the external connection. Fig. S2 (b) & (c) shows the difference in 'ideal' orientation and 'real' orientation (see Fig. 2(c)) of the MagPen prototype over the rat hippocampal slice to study micromagnetic neurostimulation. The synaptic connections between the pyramidal cell layers are also shown.

**S3. Attenuation of induced electric field and its $E_x$ and $E_y$ components with distance between MagPen Type H orientation and the neural tissue**

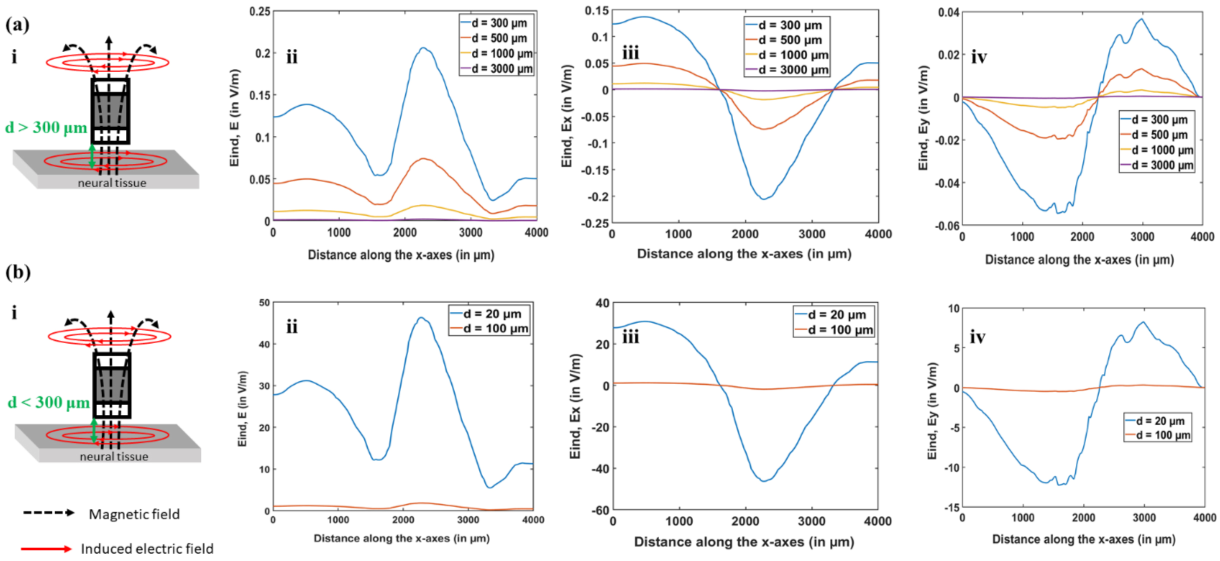


Figure S3. MagPen Type H orientation over the neural tissue. (a)- (i) For distance (d) between µcoil and neural tissue greater than 300 µm, attenuation of (ii) induced electric field (E), (iii) $E_x$ (iv) $E_y$. (b)- (i) For distance (d) between µcoil and neural tissue less than 300 µm, attenuation of (ii) induced electric field (E), (iii) $E_x$ (iv) $E_y$.

As mentioned in section 3.2 of the manuscript, in our numerical studies conducted through ANSYS-Maxwell and NEURON, we did not see the neuron eliciting an action potential beyond 300 µm distance between the neuron and the µcoil. Fig. S3 provided the detailed values for the induced electric field and its in-plane components at different distances between the µcoil and the neural tissue. Fig. S3 (a) shows how E, $E_x$ and $E_y$ attenuates at 300 µm, 500 µm, 1 mm and 3 mm distances. Fig. S3 (b) shows how E, $E_x$ and $E_y$ attenuates at 20 µm and 100 µm.

In the MagPen prototype, the µcoils are encapsulated by 2 µm thick Parylene-C (see section 2.1). This also adds to the distance between the µcoils and the neural tissue. Therefore, the use of a thickness controlled biocompatible, anti-leakage current coating of the µcoil is extremely essential. During *in vitro* EPSP recording experiments, it is very easy to exceed the 300 µm distance between the µcoils and the neural tissue and not see any biological neuron response. However, with meticulous micromanipulation of the MagPen prototype, it is not uncommon to orient the µcoil below 300 µm distance between the µcoil and the neural tissue.

**S4. Application of TTX and TTX washout protocol**

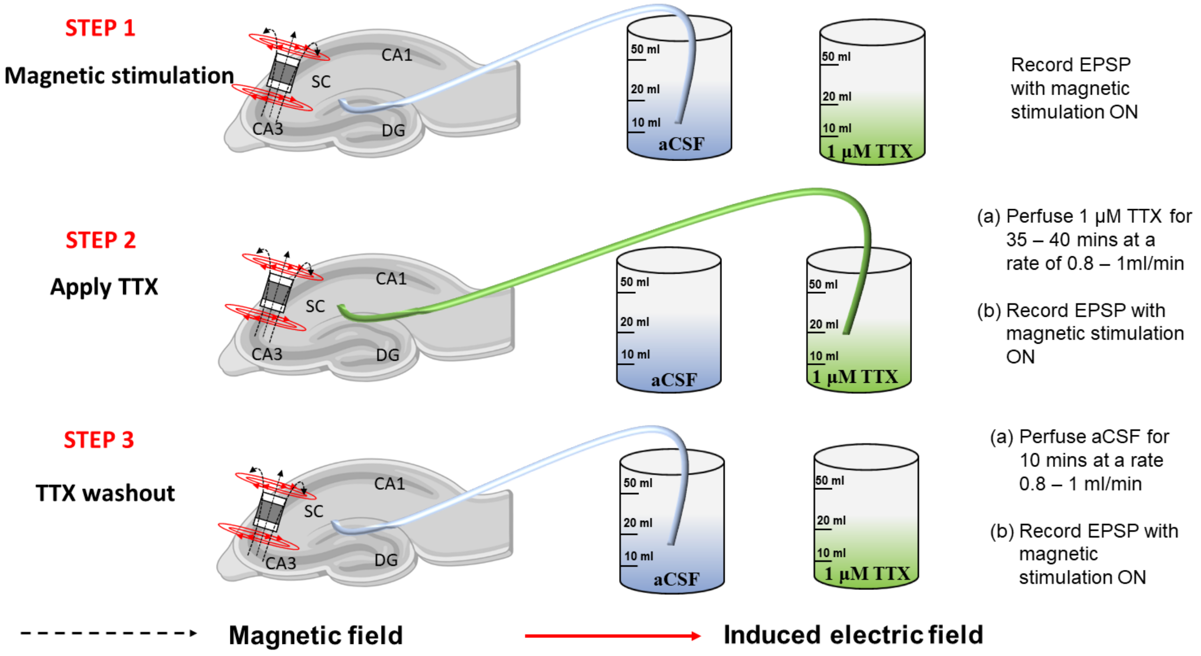


Figure S4. Pictorial demonstration of the 3-step validation process of EPSPs recorded from micromagnetic neurostimulation validation. **STEP 1: Magnetic stimulation -** Record EPSP from MagPen stimulation only. The hippocampal slice was being continually perfused by aCSF at 08-1 ml/min. **STEP 2: Apply TTX -** Perfuse tetrodotoxin (TTX) of 1 µM concentration for 35 – 40 mins at a rate of 0.8 – 1 ml/min and record EPSP stimulation from MagPen with the same parameters as in STEP 1. **STEP 3: TTX washout –** Perfuse aCSF for 10 mins at a rate of 0.8 – 1 ml/min and record EPSP stimulation from MagPen with the same parameters as in STEP 1.

The steps demonstrated in Fig. S4 are a modified version of the experimental design by Lipowsky *et al.* [1]. The comparison of the EPSPs obtained from the 3 steps are made in Fig. 7 in section 3.4.

### S5. Parameters validating successful TTX application and successful TTX washout on magnetic stimulation

Section 3.2 discusses how spatially selective and distance dependent this micromagnetic neurostimulation from this MagPen, Type H prototype is. Therefore, orienting the µcoil in the correct way and visualizing which section of the hippocampal tissue gets stimulated such that we can obtain a successful EPSP recording was extremely essential. Although these unique features for micromagnetic neurostimulation has its advantages in targeted, spatially-selective neurostimulation, yet it brings its own challenges in in vitro experiments in an academic setting.

We conducted the EPSP recordings over 6 trials out of which suppression of EPSP was successful in 4 out of 6 trials. We did not conduct TTX washout in Trial 1. The success rate for TTX washout was in 2 out of 3 trials in which suppression of EPSP due to application of TTX was observed. The peak values of the EPSPs have been calculated at a particular time instant, t ms (see Fig. S5(a)). The successful suppression of neuron response on application of TTX was quantified by equation (S1). The % drop of peak value of EPSP on application of TTX if greater than 15 %, it was a successful suppression of EPSP (see Fig. S5(b)). The average drop in 6 trials was 20 %. Greater the drop, better is the suppression. The successful TTX washout was quantified by equation (S2). The % return of peak value after washout, if less than 2 %, it was a successful washout experiment (see Fig. S5(c)). The average difference between the peak of EPSP before and after application of TTX was 0.1 %. Smaller the difference, better is the TTX washout. If the µcoils corroded during EPSP testing on hippocampal slices, new µcoils were prepared in the similar manner as discussed in section 2.1.

$$\%\ \text{drop of peak value of EPSP on application of TTX} = \left(\frac{\text{peak before TTX} - \text{peak after TTX}}{\text{peak before TTX}}\right) * 100 \quad \ldots\text{(S1)}$$

$$\%\ \textit{return}\ \text{of peak value after washout} = \left(\frac{\text{peak before TTX} - \text{peak after TTX washout}}{\text{peak before TTX}}\right) * 100 \quad \ldots\text{(S2)}$$

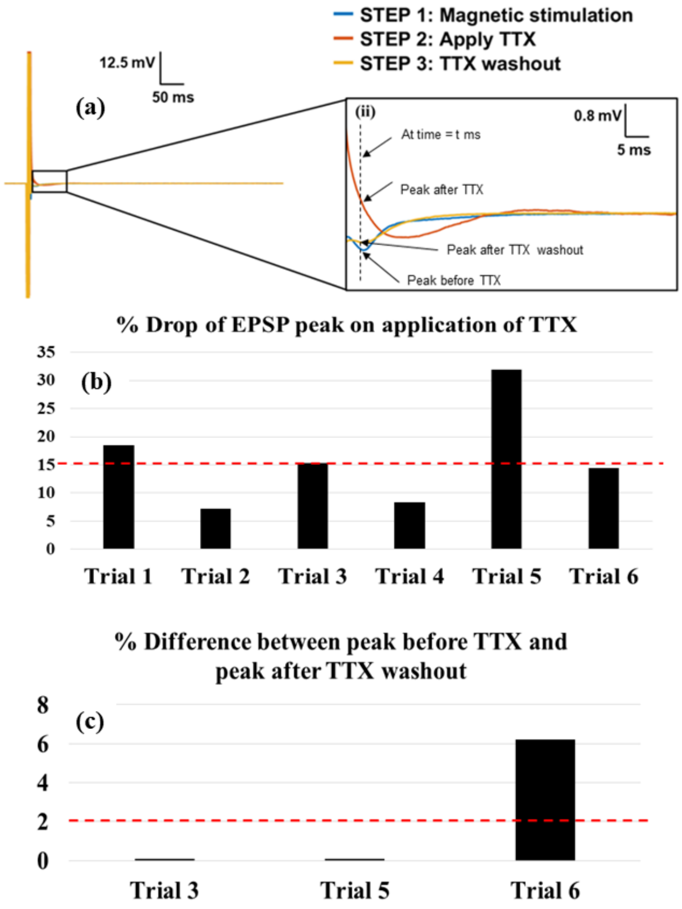


Figure S5. (a) The time instant for calculating the peaks of the EPSPs in the 3 steps. (b) % drop of peak in step 2 on application of TTX compared to step 1 and the threshold for successful suppression of EPSP marked by red dotted line. Greater the difference, better is the suppression. (c) % difference between peak value in step 1 and step 3 and the threshold for successful washout marked by red dotted line. Smaller the difference, better is the TTX washout.

**S6. Spatial components of the magnetic flux density for MagPen**

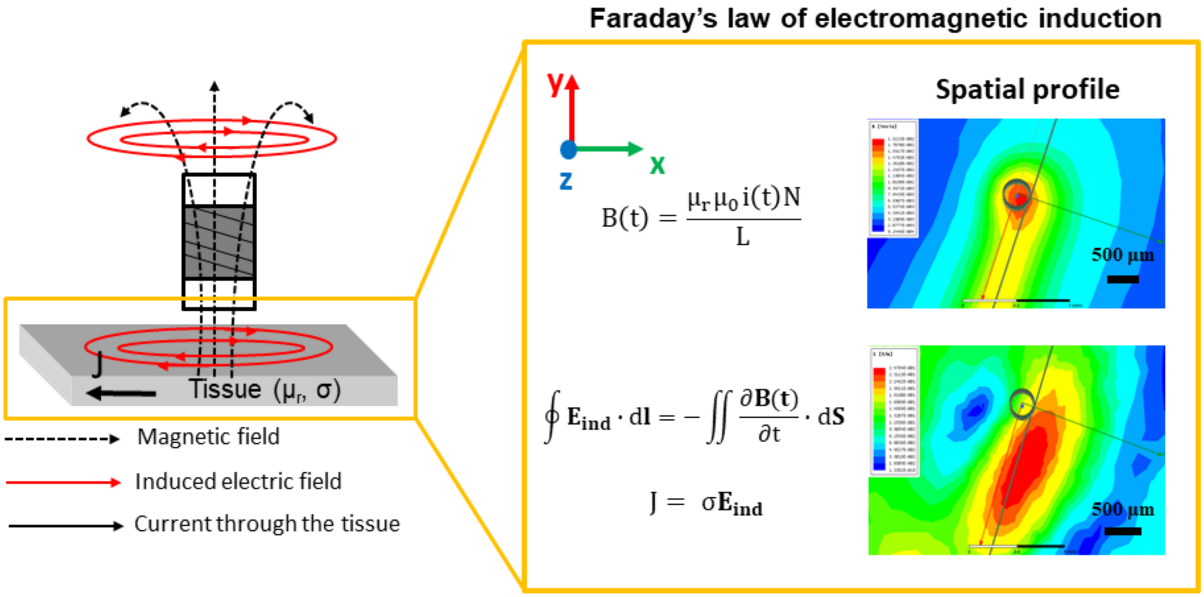


Figure S6. The spatial components of magnetic flux density, B(t) and induced electric field ($E_{ind}$) measured for the µcoil and neural tissue distance 300 µm for a sinusoidal current, i(t) of amplitude 2 A and frequency 2 kHz. Modeling performed on ANSYS-Maxwell (eddy current solver).

$\mu_r$ and $\mu_0$ are the relative permeability of the neural tissue and vacuum permeability respectively. σ is the conductivity of the neural tissue. The current that will flow through a neural tissue on stimulation by the induced electric field from these µcoils will depend on σ. When current flows through the neural tissue, the neurons start firing, i.e., it is stimulated.
